# Differential modes of orphan subunit recognition for the WRB/CAML complex

**DOI:** 10.1101/828228

**Authors:** Alison J. Inglis, Katharine R. Page, Alina Guna, Rebecca M. Voorhees

## Abstract

A large proportion of membrane proteins must be assembled into oligomeric complexes for function. How this process occurs is poorly understood, but it is clear that complex assembly must be tightly regulated to avoid accumulation of orphan subunits with potential cytotoxic effects. We interrogated assembly in mammalian cells using a model system of the WRB/CAML complex: an essential insertase for tail-anchored proteins in the endoplasmic reticulum (ER). Our data suggests that the stability of each subunit is differentially regulated. In WRB’s absence, CAML folds incorrectly, causing aberrant exposure of a hydrophobic transmembrane domain to the cytosol. When present, WRB can post-translationally correct the topology of CAML both *in vitro* and in cells. In contrast, WRB can independently fold correctly, but is still degraded in the absence of CAML. We therefore propose at least two distinct regulatory pathways for the surveillance of orphan subunits during complex assembly in the mammalian ER.

## Introduction

A large fraction of the proteome is organized into multi-subunit complexes that must be assembled at a defined stoichiometry (Huttlin et al., 2017; Marsh & Teichmann, 2015). In the cytosol, unassembled subunits expose thermodynamically unfavorable interfaces to the crowded cellular environment, which could lead to aggregation and cytotoxic effects (Sung et al., 2016; Yanagitani et al., 2017). As a result, assembly of these complexes is tightly regulated to ensure that orphan subunits, which have been synthesized in excess or cannot be assembled, are rapidly degraded to maintain cellular homeostasis (Harper & Bennett, 2016; Shemorry et al., 2013; Sung et al., 2016; Xu et al., 2016; Yanagitani et al., 2017). Despite increasing interest in cytosolic complex assembly, how multi-subunit membrane protein assembly is regulated within the endoplasmic reticulum (ER) membrane remains poorly understood (Dephoure et al., 2014).

Most membrane proteins are synthesized at the ER where their hydrophobic transmembrane domains (TMDs) must be inserted into the lipid bilayer, most commonly via the Sec61 insertion channel (Rapoport, 2007). A large proportion of membrane proteins must be further assembled into oligomeric complexes for function. Several lines of evidence suggest that this assembly process is highly regulated within the ER. First, orphan subunits of oligomeric membrane protein complexes are unstable and rapidly degraded by the ubiquitin-proteasome pathway (Juszkiewicz & Hegde, 2018; Lippincott-schwartz et al., 1988). Second, many membrane protein subunits require charged or polar residues for function or oligomerization, which prior to assembly would be exposed and thereby disfavored in the lipid bilayer. Finally, many TMDs situated at subunit interfaces are suboptimal and not predicted to insert autonomously, raising the question of how their insertion is coordinated with subunit assembly. Therefore, the mechanisms regulating oligomeric assembly within the ER are likely to be as defined and stringent as those in the cytosol.

Recent work demonstrates that in the cytosol, many multi-subunit complexes assembly co-translationally (Shiber et al., 2018): interaction between subunits occurs upon emergence of nascent domains from the ribosome, resulting in the integration of polypeptide folding and oligomeric assembly. However, unlike in the cytosol, the steric constraints of the two-dimensional lipid bilayer, combined with the fact that the Sec61 channel is surrounded by over twenty integral membrane proteins, severely limits the space available for simultaneous insertion and oligomerization. How membrane proteins overcome these additional challenges to coordinate the folding and assembly of multiprotein complexes within the ER remains unknown.

In order to better understand membrane protein assembly and quality control in the mammalian ER, we have chosen to study the regulation of the WRB/CAML complex. WRB and CAML (Get1/2 in yeast) together form an insertase for tail-anchored proteins at the ER (Vilardi et al., 2011; Vilardi et al., 2014; Yamamoto & Sakisaka, 2012). Previous work suggests that WRB and CAML stability is interdependent, consistent with it assembling into an obligate oligomeric complex (Colombo et al., 2016; Rivera-Monroy et al., 2016). The interaction between the two subunits is thought to be mediated by the TMDs, suggesting that it is likely an intramembrane signal that initiates a degradative pathway in the absence of the subunits’ cognate binding partner (Vilardi et al., 2014; Wang et al., 2014; Yamamoto & Sakisaka, 2012). Despite this, the stoichiometry of the WRB/CAML complex remains to be precisely determined, as earlier work suggests CAML is in five-fold excess of WRB *in vivo*; however, no isolated populations of CAML or WRB were detected by Blue Native-PAGE analysis of mammalian cells, suggesting CAML and WRB are always found in stable oligomeric complexes (Carvalho et al., 2019; Colombo et al., 2016).

Here we report data suggesting at least two distinct mechanisms for regulation of orphan membrane protein subunits exemplified by the WRB/CAML complex: (i) WRB is representative of a larger class of membrane subunits that insert independently but remain subject to degradation in the absence of their binding partners; (ii) in contrast CAML inserts incorrectly in the absence of WRB, aberrantly exposing a hydrophobic TMD to the cytosol, which likely acts as a flag for degradation. Upon co-expression with WRB, we observe a post-translational topological change to CAML, suggesting that WRB acts as an internal chaperone for folding and assembly of the WRB/CAML complex, consistent with assembly mechanisms observed in the cytosol. These observations set the stage for future work studying the regulation of the diversity of membrane protein subunits that must assemble at the ER.

## Results and Discussion

### WRB and CAML are destabilized in the absence of their binding partner

Earlier work has established that WRB and CAML expression is interdependent, though previous reports suggest that this regulation may occur partially at the transcriptional level (Carvalho et al., 2019; Colombo et al., 2016; Rivera-Monroy et al., 2016; Shing et al., 2017). We reasoned there may be an additional layer of regulation of WRB and CAML at the post-translational level, as has been observed for other multi-subunit complexes (Beguin et al., 1998; Bonifacino et al., 1990; Bonifacino et al., 1991; Dephoure et al., 2014; Lippincott-schwartz et al., 1988; Minami et al., 1987; Volkmar et al., 2019). To measure WRB and CAML stability, we utilized a fluorescent reporter system in which we express a GFP fusion of WRB or CAML along with RFP separated by a viral 2A sequence from a single open reading frame (Figure 1A). We first demonstrated that the introduction of these fluorescent tags does not affect WRB and CAML association in cells (Figure S1A). Therefore ratiometric analysis of the GFP:RFP fluorescence using flow cytometry can be used as a proxy for subunit stability at the protein level (Itakura et al., 2016).

**Figure 1.**
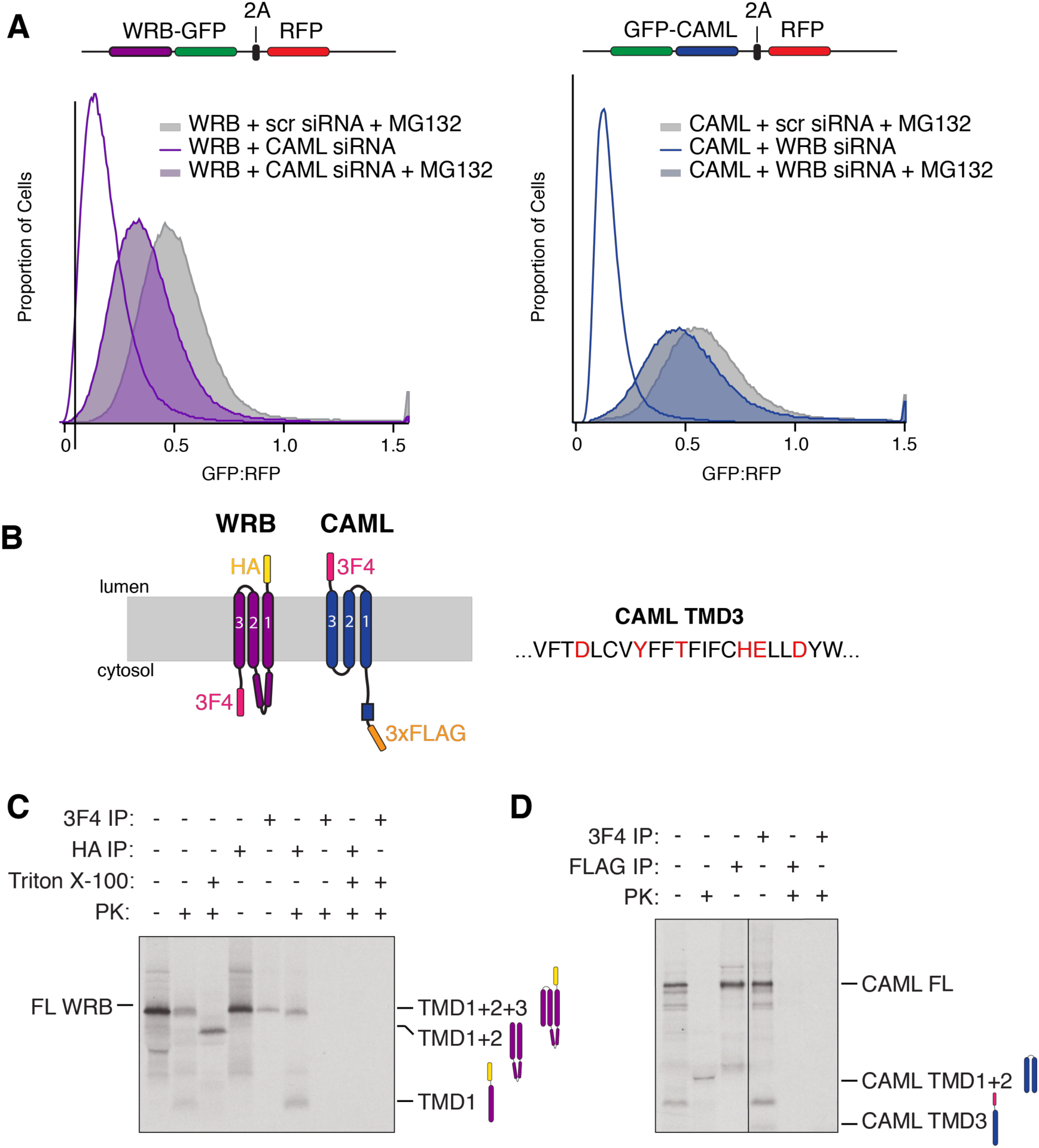
Characterization of orphaned CAML and WRB. (A) Histograms of CAML and WRB overexpression in mammalian cells as determined by flow cytometry. siRNA knockdown of their respective binding partners results in a decrease in the GFP:RFP ratio for both CAML and WRB, which is rescued by treatment with the proteasome inhibitor MG132. (B) Schematic depicting the expected topology of WRB and CAML, along with the epitope tags used for *in vitro* translation. The sequence of the third TMD of CAML is shown, with polar and charged residues highlighted. (C) ^35^S-methionine labeled HA-WRB-3F4 was translated in rabbit reticulocyte lysate (RRL) in the presence of canine-derived rough microsomes (cRM). The total products were treated with proteinase K in the presence or absence of detergent and then analyzed directly or following immunoprecipitation via the 3F4 or HA tag. WRB adopts the expected topology, with the N- and C-termini in the lumen and cytosol, respectively. The coiled-coil domain between TMD1 and TMD2 mostly protects the loop from cleavage by proteinase K, giving two major HA-tagged species in the absence of detergent. Upon the addition of detergent, the loop between TMD2 and TMD3 is cleaved, resulting in the loss of the HA tag. (D) An experiment as in (C) of FLAG-CAML-3F4. There is an untagged protease protected fragment present that likely corresponds to TMD1 and TMD2 of CAML, indicating the first two TMDs are inserted correctly. The absence of a glycosylated species despite the presence of an endogenous glycosylation site between TMD2 + TMD3 of CAML indicates the topology of this fragment is as predicted. While the first two TMDs insert properly therefore, the lack of a protease protected 3F4 fragment demonstrates that the third TMD of CAML remains aberrantly exposed in the cytosol.

Exogenous expression of either CAML or WRB individually results in rapid degradation of excess subunits, suggesting that each protein is independently unstable (approximately 80% of the CAML and 65% of the WRB that is synthesized is degraded [Figure S1B]). We observe a further decrease in the levels of both WRB and CAML upon siRNA knock down of their endogenous binding partner. This decrease in GFP:RFP ratio can be rescued using the proteasome inhibitor MG132, indicating that orphaned WRB and CAML are post-translationally regulated by the ubiquitin-proteasome pathway (Figure 1A). Consistent with tight regulation of CAML and WRB levels by the cellular quality control machinery, we observe that overexpression of either subunit results in downregulation of the endogenous protein, and upregulation of its binding partner, as has been observed for other obligate hetero-oligomeric complexes (Figure S1C; Guna et al., 2018; Juszkiewicz & Hegde, 2017).

### Two distinct mechanisms for recognition of orphan membrane subunits

Unassembled subunits in the cytosol are recognized by quality control machinery due to the aberrant exposure of thermodynamically unfavorable subunit interfaces (Yanagitani et al., 2017). However, the biophysical properties of orphan membrane protein subunits that lead to their recognition and degradation is comparatively ill defined. We therefore tested the insertion and topology of WRB and CAML to better understand how and why they are quality control substrates when unassembled.

We first demonstrated that our *in vitro* translation and insertion system, comprised of rabbit reticulocyte lysate supplemented with ER microsomes, could recapitulate the stable assembly of WRB and CAML as observed in cells (Figure S1D). We then determined the topology of individually translated CAML and WRB using a protease protection assay (Figure 1B). WRB adopts the expected topology where all three TMDs are efficiently inserted resulting in the positioning of the N- and C-termini in the lumen and cytosol, respectively (Figure 1C).

If CAML was similarly able to independently insert correctly, one would expect to observe two protected fragments: an untagged fragment representing TMDs1-2 and a 3F4-tagged protected fragment representing TMD3. However, we do not detect any 3F4-tagged protease protected species, suggesting that the C-terminus of CAML is aberrantly localized to the cytosol (Figure 1D). This observation is consistent with two possible topologies: (i) where TMDs1-2 are properly inserted but TMD3 remains in the cytosol, or (ii) where TMD1 and 3 are inserted, but TMD2 is ‘skipped’ and remains in the ER lumen. We favor the former model as the observed protected fragment is ~6 kDa in size, consistent with the expected size of TMDs1-2. Furthermore, localization of TMD2 to the ER lumen would make a naturally occurring glycosylation site between TMD2 and 3 accessible to modification; however, we do not observe a higher molecular weight glycosylated species upon translation of CAML in the presence of membranes. Therefore, the topology that best explains the protease protection data is one in which when expressed individually, only the first two TMDs of CAML insert into the lipid bilayer, while its third TMD is aberrantly exposed in the cytosol. This is consistent with the predicted inability of the third TMD to autonomously insert due to the presence of several charged and polar residues (ΔG=0.512; Hessa et al., 2007). Our biochemical evidence suggests the major population of orphan CAML is inserted in this manner, in contrast to previous reports in which both TMD2 and TMD3 are localized to the lumen, or the first and third TMD are inserted (Carvalho et al., 2019).

We next tested whether the insertion of CAML TMD3 was affected by the presence of WRB. Using a similar *in vitro* strategy, we observe that both co- and pre-expression of WRB resulted in insertion of increasing amounts of CAML TMD3 into the bilayer, as indicated by the appearance of a 3F4-tagged protease protected fragment (Figure 2A). We consistently observe an increase in insertion efficiency of TMD3 when WRB is translated prior to CAML rather than simply co-expressed, which we attribute to the additional time this provides for WRB-CAML association and TMD3 insertion.

**Figure 2.**
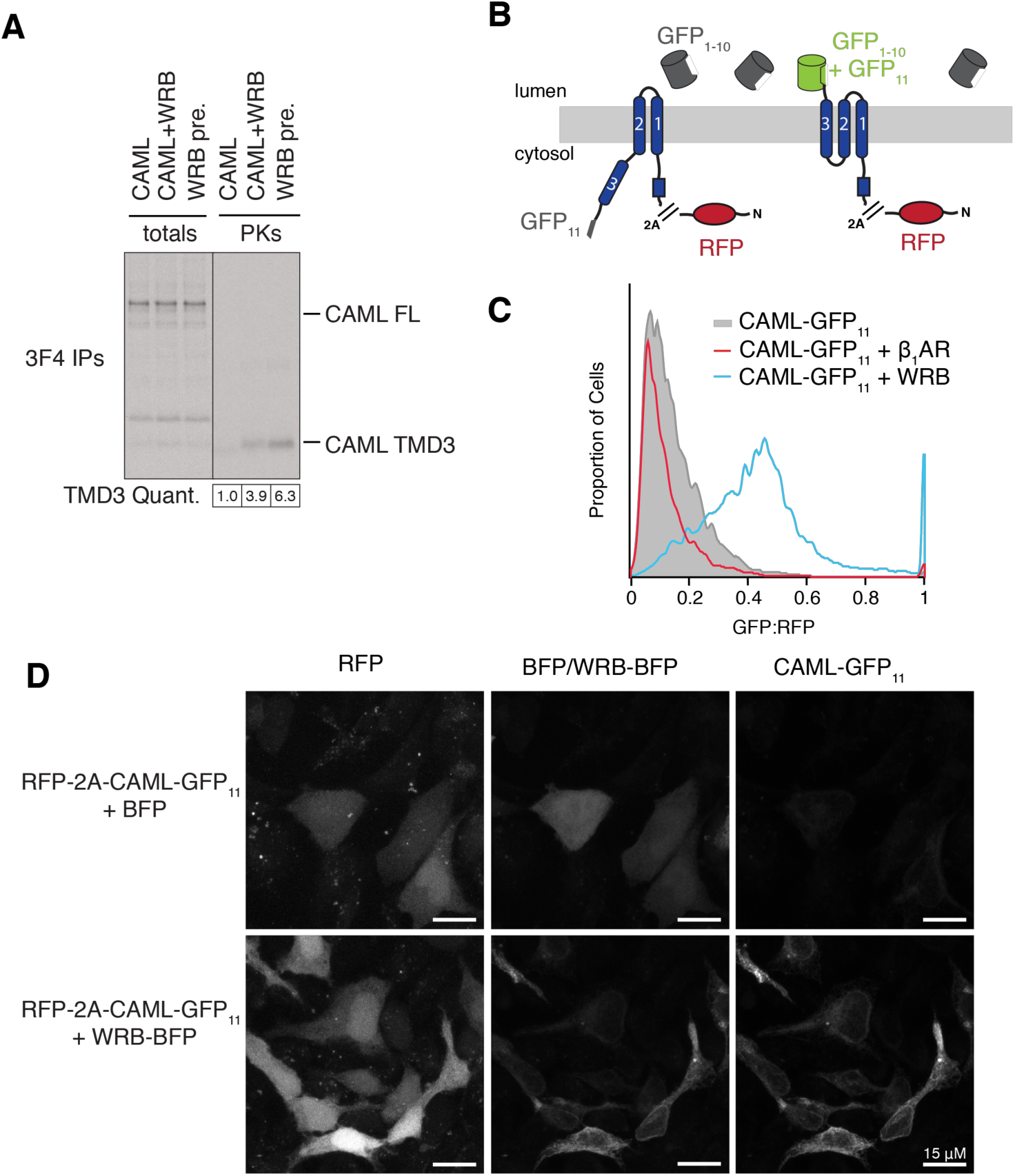
CAML requires WRB for correct insertion. (A) ^35^S-methionine labeled FLAG-CAML-3F4 was translated in RRL in the presence of cRMs either individually, alongside WRB, or with cRMs pre-loaded with WRB. Following digestion with proteinase K, total translations and digested reactions were immunoprecipitated via the 3F4 epitope tag. The positions of bands corresponding to FL CAML and CAML TMD3 are indicated. The amount of protected CAML TMD3 relative to total translated protein is indicated, normalized to the amount present for orphan CAML. (B) Schematic illustrating the split GFP system used to establish the topology of CAML in cells. CAML containing the eleventh β strand of GFP (GFP_11_) at its C-terminus was transfected into cells stably expressing the remainder of GFP (GFP_1-_10) in the ER lumen. The correct insertion of CAML TMD3 would localize GFP_11_ to the ER lumen, resulting in complementation and GFP fluorescence. (C) Flow cytometry analysis of the system described in (B) for RFP-2A-CAML-GFP_11_ expressed either alone or alongside an unrelated membrane protein (β_1_AR-BFP) or WRB-BFP. (D) ER GFP_1-10_ expressing cells were co-transfected with RFP-2A-CAML-GFP_11_ and BFP or WRB-BFP. Fixed cells were then imaged by confocal microscopy.

To confirm that WRB-dependent insertion of CAML was not an artifact of the *in vitro* system, we exploited a split GFP system to determine CAML TMD3 localization in cells (Figure 2B; Hyun et al., 2015). We generated cell lines expressing the first ten β-strands of GFP in the ER lumen. Expression of constructs that position the eleventh β-strand of GFP in the lumen, but not in the cytosol, allow for complementation and GFP fluorescence that can be measured by flow cytometry (Figure S2). When GFP_11_ is positioned at the C-terminus of CAML’s TMD3, a 5-fold increase in GFP fluorescence is observed specifically in the presence of exogenous WRB but not another unrelated membrane protein (Figure 2C). This increase in GFP fluorescence upon co-expression of CAML and WRB at the ER can be directly visualized by fluorescence microscopy (Figure 2D). The low level of GFP complementation observed when CAML-GFP_11_ is expressed individually is most likely due to partial insertion by endogenous WRB. Insertion of CAML’s TMD3 is therefore dependent on association with WRB both *in vitro* and in cells. These data are consistent with recent findings that describe a WRB-dependent conformational change to CAML in cells (Carvalho et al., 2019).

Taken together, these observations suggest that there are at least two distinct mechanisms for recognition of orphan subunits at the ER. WRB, in spite of adopting the correct topology, is destabilized in the absence of CAML. This may be due to the presence of charged or polar residues within the TMDs that would normally be shielded at its interface with CAML. Exposure of such residues could lead to recognition of unassembled WRB by membrane-embedded quality control machinery. In this way WRB is representative of a larger class of membrane protein subunits that are properly inserted and folded, yet are degraded by the ubiquitin-proteasome pathway when unassembled (Bañó-polo et al., 2017; Lippincott-schwartz et al., 1988).

Conversely, the regulation of CAML in the absence of WRB is likely at least partly due to the incorrect insertion of its TMD3. Aberrant exposure of this hydrophobic segment to the cytosol likely serves as a flag for recognition, allowing orphan CAML to exploit the cytosolic quality control machinery for degradation (Feige & Hendershot, 2013).

### CAML TMD3 insertion occurs post-translationally

Given the observation that WRB is required for insertion of TMD3 of CAML, the two most likely models are that TMD3 insertion is happening (i) co-translationally during synthesis of CAML at the Sec61 translocation channel or (ii) post-translationally, after CAML has been released from the ribosome. To discriminate between these two possibilities, we exploited our ability to pre-load membranes with either CAML or WRB to control the order of translation and insertion into the membrane (Figure S3).

One would predict that if the final folding of CAML must occur co-translationally, TMD3 insertion would be more efficient when WRB is translated first and thereby present throughout the synthesis of CAML. However, we instead find no significant difference in insertion of CAML’s TMD3 regardless of the order of WRB and CAML synthesis (Figure 3A). This is consistent with the insertion of TMD3 occurring post-translationally, after CAML and WRB have been released from the ribosome and presumably are no longer associated with Sec61. Consistent with a post-translational mechanism for insertion, we selectively recover a stable complex between CAML and WRB only after CAML has been released from the ribosome (Figure 3B).

**Figure 3.**
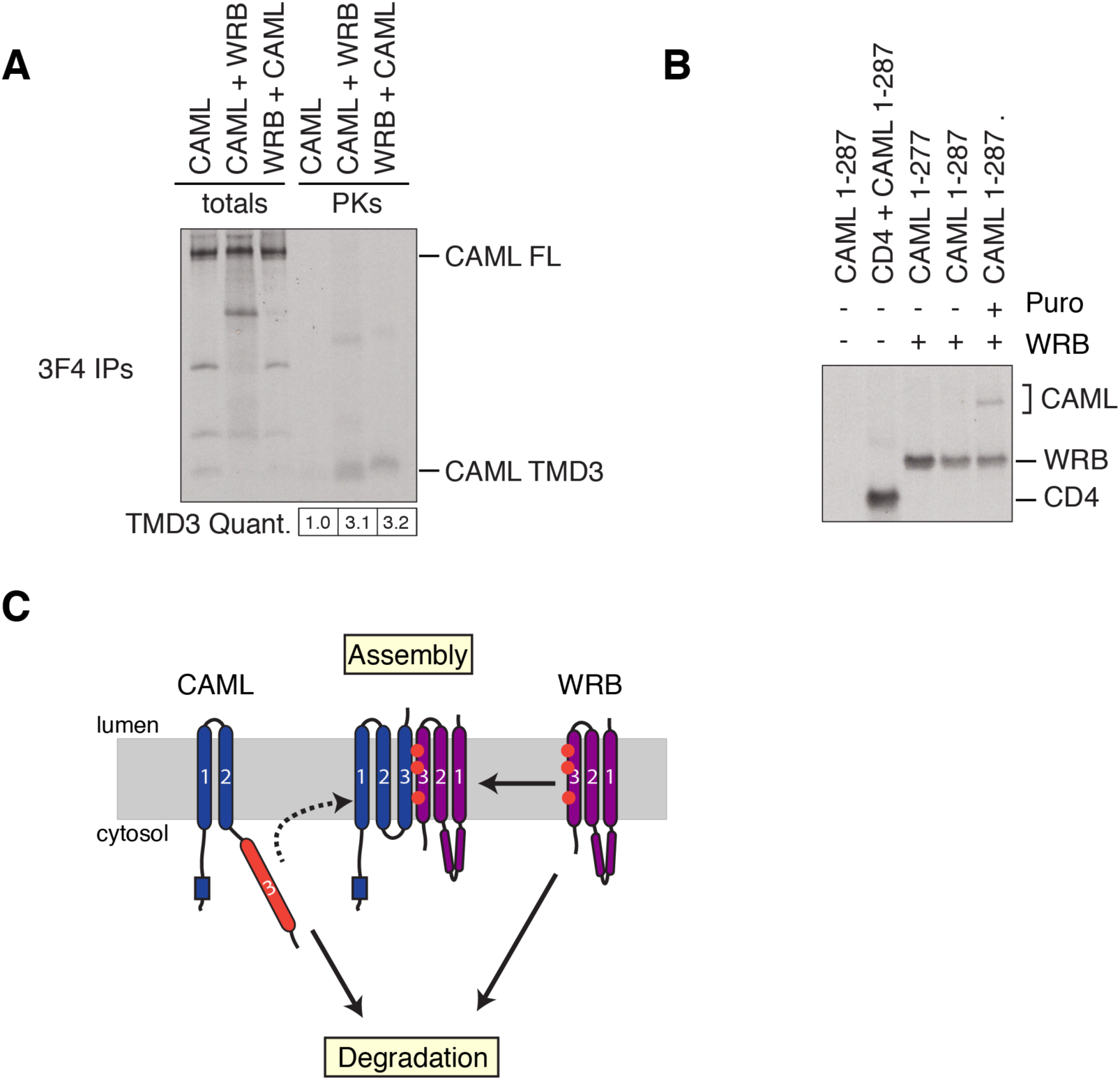
WRB causes TMD3 of CAML to insert post-translationally. (A) cRM were introduced during the translation of either i) no transcript, ii) CAML-3F4 or iii) WRB to produce i) empty, ii) ^35^S-methionine labeled CAML-3F4 or iii) ^35^S-methionine labeled WRB-preloaded membranes. Membranes were purified and nucleased to remove residual mRNA before being used in a second round of translation to produce ^35^S-methionine labeled CAML-3F4 or WRB. The insertion of TMD3 of CAML was then analyzed using a protease protection assay and immunopurification via the 3F4 tag as described in Figure 2A. The amount of protected CAML TMD3 relative to total translated protein is indicated, normalized to the amount present for orphan CAML. (B) ^35^S-methionine labeled CAML truncated at either residue 277 or 287 was produced under conditions that maintain the peptidyl-tRNA linkage. Constructs were translated in RRL in the presence of cRM preloaded with mock, an unrelated membrane protein (CD4) or FLAG-WRB (labeled with ^35^S-methionine). Puromycin treatment was used to release the nascent protein from the ribosome, and then the membranes were solubilized with digitonin before affinity purification under native conditions via the FLAG tag of WRB. The position of bands corresponding to CD4, WRB and CAML is shown. (C) A proposed model for the regulation of assembly of the WRB/CAML complex: upon initial synthesis CAML is misfolded, aberrantly localizing TMD3 to the cytosol. Only after post-translational interaction with WRB does CAML insert and fold correctly. For simplicity we have depicted a single WRB/CAML heterodimeric interaction, but WRB may operate catalytically to fold multiple CAML subunits to account for the observed excess of CAML relative to WRB (Colombo et al., 2016). In the absence of WRB, TMD3 likely serves as a flag for degradation of orphaned CAML, which can exploit the cytosolic quality control machinery for recognition and degradation. In contrast, WRB independently adopts the correct topology upon synthesis, yet is robustly degraded in the absence of CAML. Together WRB and CAML therefore represent two distinct mechanisms for stoichiometric regulation within the ER membrane.

Together this suggests a working model for the folding and assembly of the WRB/CAML complex (Figure 3C). Initial recruitment of WRB occurs post-translationally, and is likely mediated by the first two TMDs of CAML, which insert independently. Whether this partially folded version of CAML must be stabilized by either an intramembrane and/or cytosolic chaperone prior to association with WRB remains to be determined. Similarly, unassembled WRB may also require stabilization by a membrane-embedded chaperone, to provide sufficient time for association with CAML. Upon binding, WRB is able to catalyze the insertion of CAML’s TMD3 into the ER membrane, thereby acting as an internal chaperone for the folding and assembly of the WRB/CAML complex. This strategy allows insertion of this poorly hydrophobic TMD, which is not independently recognized by Sec61. The lack of certainty surrounding the complex stoichiometry means that we cannot conclude whether WRB is acting on a single CAML subunit as part of a stable complex or whether it is acting catalytically on multiple copies of CAML. Given WRB/CAML is itself a membrane protein insertase, it is possible that this post-translational insertion is a unique feature of assembly of this complex. However, evidence for other such post-translational topological changes in polytopic proteins suggest the possibility that it could be a more general mechanism utilized by multi-subunit complexes (Hegde & Lingappa, 1999; Lu et al., 2000; Serdiuk et al., 2016).

However, if either CAML or WRB cannot assemble, their orphan forms are recognized and degraded by the ubiquitin-proteasome pathway. This recognition occurs via two distinct mechanisms: (i) improperly folded CAML aberrantly exposes its TMD3 to the cytosol, likely making it a target for the cytosolic quality control machinery, while (ii) WRB, though folded correctly, must be recognized due to the aberrant exposure of its subunit interface within the lipid bilayer. As eukaryotic membrane protein subunits differ enormously in size, topology, and the biophysical properties of their exposed interfaces, interaction with such a diverse range of substrates would require a network of chaperones in the ER membrane that remain to be identified. This work therefore sets the stage for future research to determine both the triage factors that target unassembled proteins towards either a biosynthetic or degradative fate, and how these pathways are coordinated to ensure the precise assembly of multi-subunit complexes at the ER.

## Materials and Methods

### Plasmids, antibodies, siRNAs, and purifications

Constructs for expression in cultured mammalian cells were generated in either the pcDNA5/FRT/TO (Thermo Scientific) or pcDNA3.1 backbone. To create the fluorescent reporters described in Figure 1A, cDNA for human CAML and WRB was purchased from IDT and inserted into a pcDNA5 vector expressing GFP-2A-RFP resulting in an N-(CAML) or C-terminal (WRB) GFP fusion. In order to express the split GFP_1-10_ in the ER lumen, a construct expressing the human calreticulin signal sequence preceding a GFP_1-10_-KDEL was also generated in pcDNA5 (Cabantous et al., 2004; Kamiyama et al., 2016). WRB-BFP, the turkey β_1_-adrenergic receptor, CAML-GFP_11_ (GFP_11_ tag: RDHMVLHEYVNAAGIT), cytosolic RFP-2A-GFP_11_, and RFP-2A-VAMP-GFP_11_ were inserted into pcDNA3.1 for transient mammalian expression. All experiments were performed in the Flp-In T-REx 293 cell line (Thermo Scientific). The mCherry and mEGFP versions of RFP and GFP are used throughout this manuscript, though are referred to as RFP and GFP for simplicity in the text and figures.

Constructs for expression in rabbit reticulocyte lysate (RRL) were based on the SP64 vector (Promega). For all protease protection assays (Figures 1C, 1D and 2A and 3A) CAML was expressed with an N-terminal 3xFLAG tag and a C-terminal 3F4-tag (Stefanovic & Hegde, 2007) while WRB was appended with an N-terminal 1xHA tag and C-terminal 3xFLAG tag. Tags were chosen to minimize interference with TMD insertion, with those containing multiple charged or polar residues being placed on the cytosolic face.

Purification of GFP-tagged CAML and WRB from mammalian cells were performed using an anti-GFP nanobody (Kirchhofer et al., 2009; Pleiner et al., 2015). Briefly, cell lines of GFP-2A-RFP, WRB-GFP-2A-RFP, and GFP-CAML-2A-RFP were cultured in 10 cm dishes until 70% confluent, induced with 1 µg/mL doxycycline and harvested after 24 hours. Cells were lysed in Solubilization Buffer (50 mM HEPES pH 7.5, 200 mM KOAc, 2 mM MgOAc_2_, 1% Digitonin, 1X protease inhibitors, 1 mM DTT) for 20 minutes at 4 °C. Pierce Streptavidin Magnetic Beads (Thermo Scientific, 88817) were equilibrated with 3.75 µg biotinylated anti-GFP nanobody in Wash Buffer (50 mM HEPES pH 7.5, 100 mM KOAc, 2 mM MgOAc_2_, 0.25% Digitonin, 1 mM DTT). Cell lysate was incubated with anti-GFP nanobody immobilized on Streptavidin support for one hour at 4 °C. GFP-tagged proteins were eluted with 0.5 µM SUMOstar protease and used directly for Western blot analysis.

Antibodies were purchased against CAML (Synaptic Systems, 359 002), WRB (Synaptic Systems, 324 002), and α-tubulin (Sigma, T9026). The antibody against 3F4 was a gift from the Hegde lab and has been previously described (Chakrabarti & Hegde, 2009). Secondary antibodies used were HRP-conjugated Goat Anti-Rabbit (BioRad, 170-6515) and Anti-Mouse (BioRad, 172-1011). Anti-FLAG (A2220) and HA resin (A2095) were obtained from Sigma (St. Louis, MO). Pre-designed Silencer Select siRNA from Thermo Fisher were obtained for CAML (s23760, s2371, s2372) and WRB (s14904, s14905).

### Mammalian *in vitro* translation

Translation extracts were prepared using nucleased rabbit reticulocyte lysate (RRL) and canine pancreatic microsomes (cRMs) as previously described (Sharma et al., 2010; Walter & Blobel, 1983). Briefly, templates for *in vitro* transcription were generated by PCR using primers that included the SP6 promoter at the 5’ end and a stop codon followed by a short untranslated region at the 3’ end. In the case of Figure 3B, primers were designed to anneal upstream of the stop codon in order to generate a truncated protein product in which the C-terminal residue is a valine, known to stabilize the peptidyl-tRNA product (Shao et al., 2013). Transcription reactions were incubated at 37 °C for 1 hour, and then used directly in a translation reaction, which was incubated for 35 minutes at 32 °C.

To generate pre-loaded membranes of either WRB or CAML, as used in Figures 2A and 3A, non-nucleased cRMs were included in an initial translation reaction for 20 minutes with mRNA for the desired protein. Membranes were purified by pelleting for 20 minutes at 55,000 rpm in a TLA55 at 4 °C through a 20% sucrose cushion in physiological salt buffer (50 mM HEPES pH 7.5, 100 mM KOAc, 2 mM MgOAc_2_). Pellets were resuspended in physiological salt buffer at a concentration of A_280_ ~80. Pre-loaded membranes were nucleased as previously described: after adjusting to 1 mM Ca^2+^, membranes are incubated for 7 minutes at 25 °C with the Ca^2+^ activated nuclease from *S. aureus*, which is then quenched by addition of 2 mM EGTA pH 8.0 (Pelham & Jackson, 1976). Nucleased membranes were either used directly in a second translation/insertion reaction or aliquoted and flash frozen for storage at −80 °C. We saw no reduction in translation and insertion efficiency after freezing.

Protease digestions were performed on ice by addition of 0.5 mg/mL proteinase K to translation reactions and incubated for 1 hour. The digestion was quenched by addition of 5 mM PMSF in DMSO, followed by transfer to boiling 1% SDS in 0.1 M Tris pH 8.0 (RT). Immunoprecipitation of protected fragments was performed in IP buffer (50 mM HEPES pH 7.5, 100 mM KOAc, 2 mM MgOAc_2_, and 1% Triton X-100).

Co-immunoprecipitation experiments (Figures 3B and S1D) were performed by setting up translation reactions in the presence of cRMs, and then purifying the membranes via pelleting for 20 minutes at 55,000 rpm in a TLA55 at 4 °C through a 20% sucrose cushion in physiological salt buffer. The pellets were resuspended in physiological salt buffer before solubilization of the membranes in 1% digitonin. The samples were then diluted four-fold and immunoprecipitated with anti-FLAG resin.

### Cell culture

Stable cell lines expressing GFP-CAML-2A-RFP, WRB-GFP-2A-RFP, or ER GFP_1-10_ were generated using the Flp-In T-Rex 293 Cell Line (Thermo Scientific) according to the manufacturer’s instructions. In brief, a 10 cm dish of cells was transfected with 9 µg of Flp-Recombinase (plasmid pOG44) and 1 µg of a specific pcDNA5/FRT plasmid using TransIT-293 transfection reagent (Mirus, MIR2705). 48 hours after transfection, cells were selected with 100 µg/mL hygromycin in DMEM media containing 10% fetal bovine serum and 15 µg/mL blasticidin. After 7-10 days the resulting isogenic cell population was expanded for maintenance and preservation.

All siRNA experiments (Figure 1A) were performed in a 6-well tissue culture plate. Cells were transfected with 3 ng of siRNA per well using RNAiMAX lipofectamine (ThermoFisher, 13778150). After 48 hours, the integrated reporter gene was induced with 1 µg/mL doxycycline for 24 hours. Before analysis, cells were treated with either 12 µM of the proteasome inhibitor MG132 (Calbiochem, 474790) or a DMSO control for 8 hours. Live cells were first incubated with trypsin before collection, pelleted, and resuspended in 300 µL of PBS containing 1 µM Sytox Blue Dead Cell Stain (ThermoFisher, S34857) and analyzed on a Miltenyi Biotech MACSQuant VYB Flow Cytometer. Data analysis for all flow cytometry experiments was performed using the FloJo software package.

GFP complementation experiments by flow cytometry were also performed in a 6-well tissue culture plate. Expression of the GFP_1-10_ protein was induced for 72 hours with 100 ng/mL doxycycline before transfection of 0.17 µg of GFP_11_ constructs, 0.17 µg of WRB-BFP or β_1_AR-BFP, and 1.36 µg of pcDNA3.1 backbone with TransIT-293 transfection reagent. Cells were harvested and analyzed by flow cytometry 24 hours after transfection. For analysis by confocal microscopy, the cells were grown in a 24-well tissue culture plate containing 12 mm glass coverslips coated in poly-D-lysine. The induction and transfection conditions for imaged samples were identical as those subjected to flow cytometry, except cells were transfected with 30 ng of RFP-2A-CAML-GFP_11_, 30 ng of BFP or WRB-BFP and 240 ng of pcDNA3.1 backbone. The cells were fixed for fluorescence microscopy according to standard protocol. In brief, the cells were washed with PBS before being incubated with 3.6% paraformaldehyde for 30 minutes. The cells were washed again, treated with Prolong Diamond Antifade Mountant (ThermoFisher, P36961) and sealed onto a slide. Imaging was performed using an LSM 800 confocal microscope (Zeiss).

For overexpression of GFP-tagged CAML and WRB (Figure S1B), cells were cultured in 6-well tissue culture plates, induced with 1 µg/mL doxycycline for 24 to 72 hours, and harvested in 5 mM EDTA pH 8.0 in 1X PBS. Cells were lysed with NETN lysis buffer (250 mM NaCl, 5 mM EDTA pH 8.0, 50 mM Tris-HCl pH 8.0, 0.5% IGEPAL CA-630, 1X protease inhibitors) for 1 hour at 4 °C. Cell lysates were used directly for analysis by Western blot. Samples were normalized by cell counting prior to lysis.

## Acknowledgements

We thank T. Pleiner for help with GFP affinity purification, and the Caltech Flow Cytometry facility for their help with FACS experiments. Confocal imaging was performed in the Caltech Biological Imaging Facility, with the support of the Caltech Beckman Institute and the Arnold and Mabel Beckman Foundation. The antibody against 3F4 was a kind gift from Ramanujan S. Hedge. This work was supported by the Heritage Medical Research Institute, the Kinship Foundation, and the Pew-Stewart Foundation.

---

**Figure S1.**
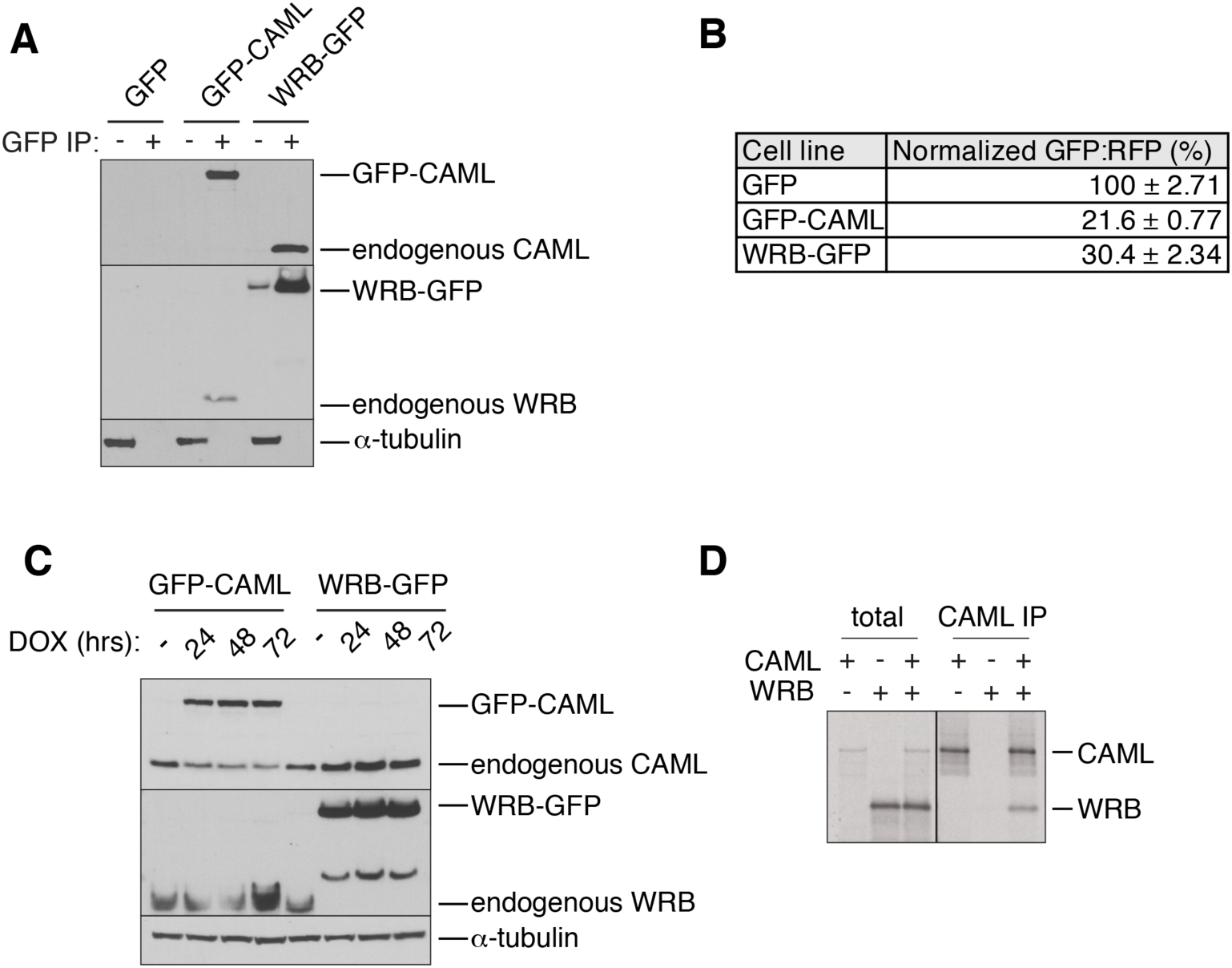
WRB and CAML interact *in vivo* and *in vitro*. (A) Detergent solubilized lysates from stable cell lines expressing GFP, GFP-CAML, or WRB-GFP were affinity purified under native conditions via the GFP epitope. Samples were analyzed for co-purification of endogenous CAML and WRB with the fluorescently-labeled subunits. (B) Exogenous expression of GFP-CAML or WRB-GFP was induced with doxycycline (DOX) for 24, 48 and 72 hours before analysis by Western blot with antibodies against CAML, WRB and α-tubulin. (C) Normalized GFP:RFP ratios from flow cytometry analysis of stable cell lines expressing GFP-2A-RFP, GFP-CAML-2A-RFP, or WRB-GFP-2A-RFP. The GFP:RFP ratios were measured in triplicate, and normalized to the GFP-2A-RFP cell line. Displayed are the means and three standard deviations. (D) ^35^S-methionine labeled FLAG-CAML or WRB were translated alone or in combination in RRL in the presence of cRMs. Following purification of the resulting microsomes and detergent solubilization, samples were affinity purified via the FLAG epitope for analysis.

**Figure S2.**
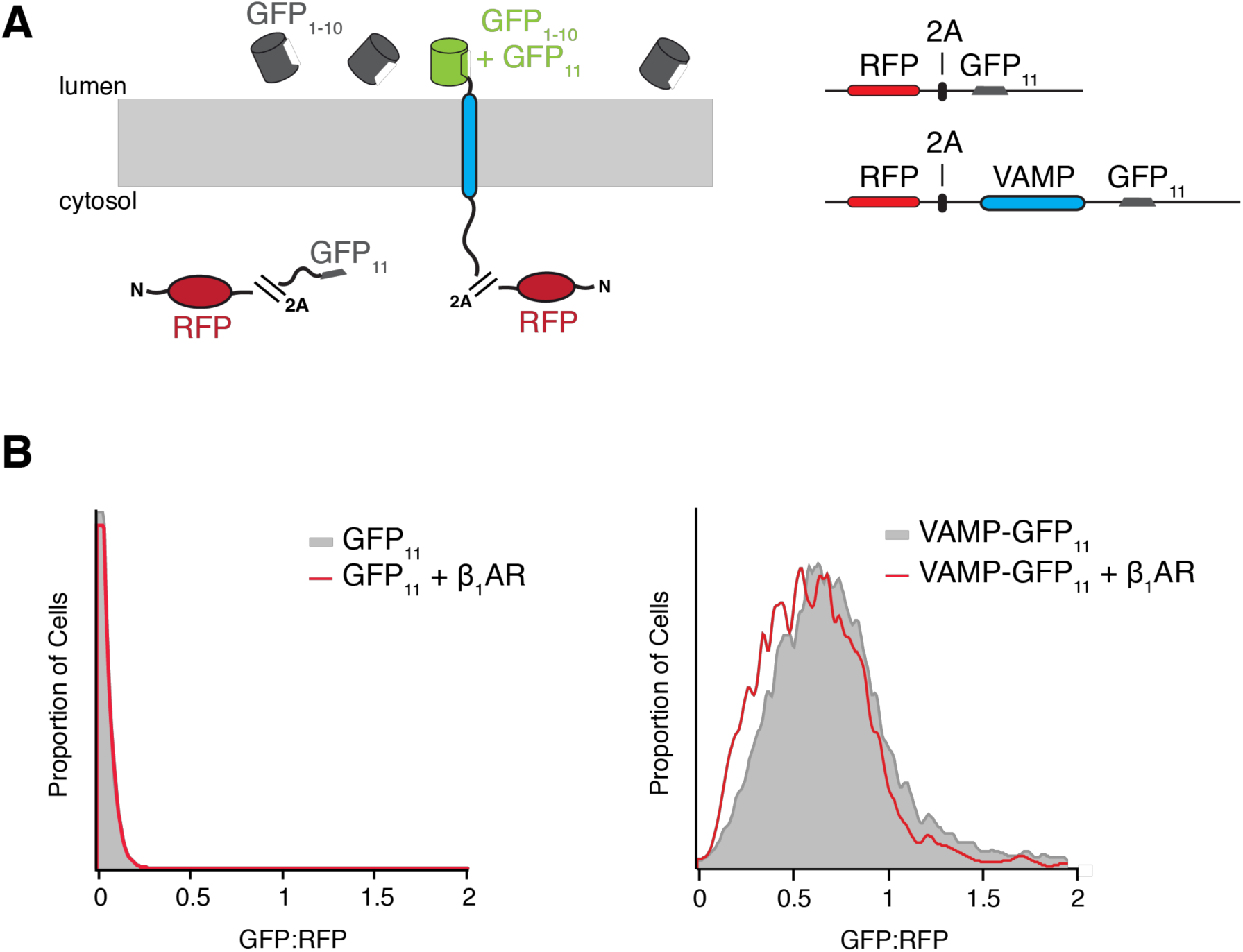
Characterization of the split-GFP system to investigate membrane protein topology in cells. (A) The first ten β-strands of GFP (GFP_1-10_) are stably expressed in the ER lumen. An increase in GFP fluorescence is detected upon complementation with the eleventh β-strand of GFP (GFP_11_) when it is localized to the lumen. In order to characterize the system, we demonstrate that expressing a cytosolic GFP_11_ results in no increase in GFP fluorescence. However, when GFP_11_ is localized to the ER lumen, such as when affixed to the C-terminus of the tail-anchored protein VAMP, an increase in GFP fluorescence is observed. In order to correct for relative expression of the cytosolic and lumenal GFP11 constructs, they are expressed as part of an open reading frame containing contain RFP separated by the viral 2A sequence. (B) Flow cytometry analysis of the system described in (A) of RFP-2A-GFP_11_ or RFP-2A-VAMP-GFP_11_ alone or in the presence of an unrelated membrane protein β_1_AR-BFP. The GFP:RFP ratio was measured by flow cytometry.

**Figure S3.**
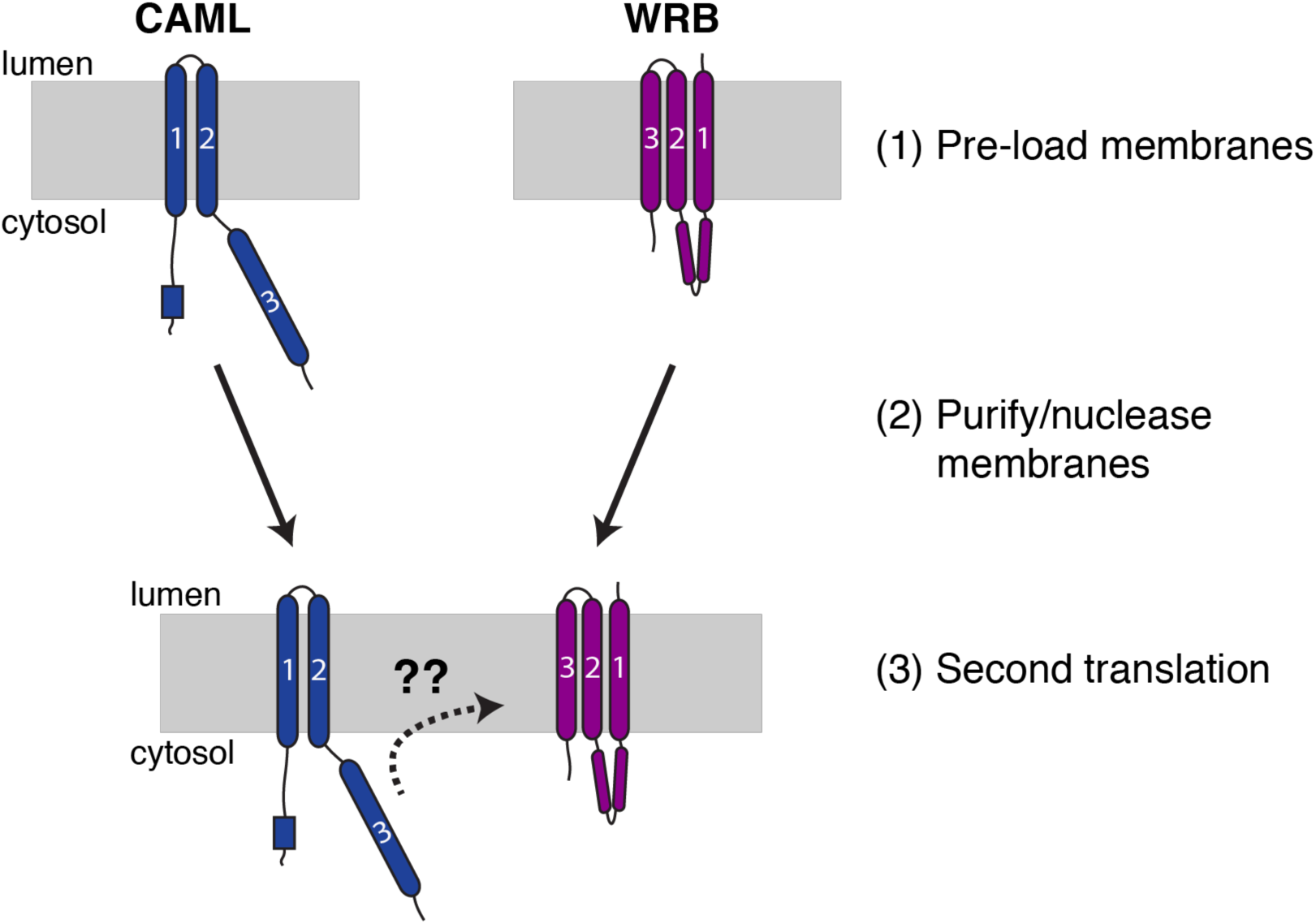
Schematic showing cRM preloading to determine the effect of biasing the order of subunit translation. After an initial incubation to allow translation and insertion of each subunit, membranes which now contain a radioactively labeled and inserted subunit are re-purified. To precisely limit the synthesis of each subunit to the pre-loading step and prevent purification of the mRNA along with the membranes, we also include a nuclease step after the purification. These membranes can then be used in a subsequent translation reaction, where the second subunit is introduced.

